## Supplemental Data for "PHERI - Phage Host Exploration pipeline"

### Supplement

#### Details of pipeline execution

Our pipeline consists of python scripts and publicly available bioinformatics software. We have taken care of the code's readability, sustainability and extensibility. We implemented our pipeline in workflow management system Snakemake [1] that was primarily designed for writing reproducible bioinformatics pipelines. Snakemake is inspired by GNU make, but it uses python-like syntax with elements similar to pseudo code. Furthermore, it is fully portable, depending only on Python executables and libraries. When the Snakemake is executed, it runs the first rule in the specified Snakefile. If the rule has missing input files, it scans through the whole Snakefile and looks for rules that are capable of creating required files. This process is repeated until there is a rule which can be completed or until there is a rule whose input is not possible to create by any other rule. In the former case the execution starts running, in the latter case an error message is displayed. By this approach it is ensured that we do not execute any unnecessary rules nor any rules that have been already completed. This is an important feature for our program as some rules can take several hours to complete, even on a powerful computational cluster. Another useful characteristic of the Snakemake engine is the ability to produce graphical visualization of a particular Snakefile in the format of a directed acyclic graph.

The pipeline starts with downloading of publicly available data. After downloading, we merge all records and eliminate duplicate records. Next, we search for genes in sequences of the bacteriophages. Consequently, phage genomes represented as sets of genes are split into a training and a testing set. Similarity between the genes from the training set are calculated and based on those, clusters of similar genes are produced. From these clusters, the binary matrix is created. This matrix is later used in classification.

### Downloading phage genomes

The first step in our pipeline is downloading data from three publicly available databases. Although they cover the majority of currently sequenced and published phages, we made this step easily extensible for adding new sources of information in the future.

#### GenBank database

National Center for Biotechnology Information (NCBI) provides access to the GenBank [2] database. This database is a comprehensive source of genomic data with more than 200 million genomic sequences of all life's domains. NCBI administers the GenBank database free of charge and gives researchers the possibility to access data through various interfaces as web-based retrieval services, FTP and Entrez [3]. Despite these facts, there are shortcomings of using GenBank. With the recent breakthrough of high-throughput medical technologies the amount of data flowing into the GenBank database every day is enormous. Therefore, it is unreasonable to check all the data. Sequences are primarily submitted by individuals from all around the globe and are not thoroughly reviewed. This causes redundancy of sequences and sometimes it even creates contradictions between information in the system. We obtained data using python library Biopython [4], which implements python wrapper NCBI Entrez. Besides that, we used Biopython to facilitate processing of standard file formats used in bioinformatics. When downloading sequences, we also created unique identifiers for each record. Those were used later in the pipeline. Reasons behind the decision to use custom identifiers was the ability to remove duplicated sequences and the possibility to find out multiple sources of each sequence in our dataset. Downloading from GenBank was our largest source of data with 6091 downloaded records.

#### ViralZone database

ViralZone provides highly reliable data about viruses, including bacteriophages. Information about the structure of a capsid, a genome, life cycle, replication mechanisms, taxonomy, geographical location and host are included. This website does not store sequences internally, rather it delivers links to the RefSeq [4, 5] database. Compared to GenBank, RefSeq database contains fewer sequences, but all of these sequences are curated and manually reviewed.

Our custom script was used to download records from the RefSeq database. Although, large portion of sequences downloaded was identical with GenBank records, some sequences were unique. Another advantage in performing this action was that it enabled us to pair genomic sequences from RefSeq with more comprehensive information from ViralZone portal. By performing this process, we obtained 2070 records.

#### PhagesDB

PhagesDB is a database specialized in bacteriophages infecting bacteria from phylum Actinobacteria. This phylum is of great importance, because of its contribution to the soil system in the form of decomposing of organic matter. This phylum also contains the genus Mycobacterium, which includes pathogens causing tuberculosis and leprosy in humans [6]. The database was designed to avoid the time between sequencing and data availability. Authors declare, at the time of their publication, there were more than 600 records of bacteriophages that were not yet in GenBank. Furthermore, PhagesDB stores more biologically relevant data, such as discovery details, sequencing details, characterization details, sequence file and a plaque picture. We downloaded 2567 phage records with our automatized script using the publicly available Application Programming Interface (API).

#### Merging and removing of duplicate records

Downloaded records were highly redundant, mainly because many of those records were present in more databases at once. To solve this issue, we merged datasets together and removed duplicate records. For merging we used the standard unix command **cat**. For removing duplicated sequences, we created a custom Python script. This script made use of custom identifiers, which were assigned to every sequence that was downloaded. In case more identical sequences were found, their custom identifiers were rewritten with the identifier of the first sequence. This approach preserved the relationships between one particular sequence and all data related to it. Thus, we were able to track phages, based on their identifiers, to their source databases and also connect them with all data that was already downloaded. After removal of the duplicated sequences, our dataset contained 7064 phage records. This suggests a high duplication ratio between the databases.

#### Extraction and annotation of genes

We used a publicly available pipeline called Prokka [7] to identify and annotate genes. First, coordinates of coding DNA sequences (CDS) were found with Prodigal tool [8]. After the locations of genes are predicted, Prokka can start to annotate functions of all CDSs. This is usually done through comparison of a sequence to a database of sequences with an experimentally determined function. The function of protein with the best match is then assigned to the new CDS. Prokka, by using this approach, searches through multiple databases. Starting with the most reliable source, which is usually the smallest, it scans all the databases, continuing with the less accurate one. The databases used with their corresponding order are as follows: An optional user-defined database, UniProt [9], RefSeq [5], Pfam [5, 10] and TIGRFAM [11]. If no match is found across databases, protein is labelled as *hypothetical* *protein*. After annotation, resulting genes were selected and saved to files in format suitable for further use in the pipeline.

#### Datasets used

At the beginning of this step, our dataset consisted of: phage genomic records, their corresponding genes, information about phage hosts and functional annotation of genes. As our work used techniques of supervised machine learning, we needed to split the dataset to a training set and a testing set. Furthermore, we created a set for other records, which we decided not to use due to missing information about hosts or due to hosts outside of our group of interest. We decided to group phages according to the genus of their hosts. After calculating the number of phages in each group, we selected the first 50 genera with the highest count of records as groups of our interest. All other genera were excluded from the dataset due to the insufficient number of samples. Phages without information about their hosts were also excluded. Subsequently, we divided remaining data into a training set and a testing set at a ratio of 4:1. The resulting training set included 4723 records of bacteriophages and the resulting testing set included 1202 records.

#### Alignment of genes

In our pipeline, we used alignment to find similarity scores between genes in the training set. We needed these scores later in the pipeline at the clustering step. Software CrocoBLAST [11, 12] was used for this purpose. CrocoBLAST is a wrapper around BLAST algorithm which makes better use of parallelization than standard BLAST maintained by NCBI. With this software we were able to reduce the time needed for the alignment step from around four days to one day. Resulting file was in tab separated format, where first column was gene identifier of query sequence, second column was gene identifier of the target sequence and third column was e-value of alignment.

#### Clustering

Firstly, we used the Markov Cluster Algorithm implemented in package MCL [13]. This software is popular in the bioinformatics community for its capabilities to work with big data. As input data we used E-values from CrocoBLAST results. These E-values were automatically transformed into a similarity score to enable creation of the adjacency matrix. We used the value 1.2 as the inflation parameter. This was the smallest value recommended by the developers. The reason for using the smallest possible inflation value was that we wanted to create clusters that are as large as possible. This approach also reduced the number of different clusters. In our work, we needed as few reasonable clusters as possible, mainly due to the number of phage records in our dataset. If we had too many clusters, we would have too many features for the classifier and we would risk overfitting of our final models to the training dataset. We created 32281 gene clusters.To get more accurate clustering, we tried a different approach, where we determined similarity score from the global alignment. All the matches from CrocoBLAST search were aligned with the needleman-wunsch algorithm [14] and the resulting score of alignment was used instead of the E-value. Implementation of the Markov Cluster Algorithm was executed with an inflation parameter value of 1.2, without any transformation of the similarity score. By this approach we created 17223 final clusters.

In this work, we also experimented with a different clustering algorithm, Spectral clustering. SCPS implementation [15] was tested in the pipeline. Although authors of this software declare the quality of clusters quantified by a measure that combines sensitivity and specificity to be better by 28% in comparison to MCL algorithm, our memory was not sufficient for the program to run with all the input data.

#### Annotation of gene clusters

Clusters of genes were annotated to determine their function. We expected proteins with similar biological function to be included in the same cluster. For functional annotation of particular proteins we used software InterProScan [16]. The reason to use InterProScan annotations instead of Prokka annotations was because InterProScan annotation had better standardized descriptions of functions. This tool scans given protein sequences against the protein signatures in databases PROSITE [17], PRINTS [18], Pfam [10], ProDom [19] and SMART [20]. After acquisition of annotations for all proteins in a particular cluster, we calculated the number of occurrences of each distinct biological function. These statistics represented our annotation of a certain cluster. We manually reviewed the annotations of the biggest clusters to evaluate the relevance of created clusters. Although a lot of proteins remained without any assigned function, our expectation of clusters containing proteins with similar function was met in most cases. Therefore, we assumed that reasonable clustering was achieved.

#### Reducing phage genomes

One of the most crucial parts of our analysis was the matrix created in this step. Rows in this matrix represented particular phage records and columns represented particular protein clusters. The entry $a_{i,j}$ in the matrix was filled with a number of genes from cluster j belonging to phage i. Custom python script and files produced in previous steps were used for this task. Resulting matrix contained 4723 rows and 32281 columns and served as a main input file for the machine learning algorithms.

#### Principal component analysis

Principal component analysis (PCA) is a method mostly used for visualization of high dimensional data. For a matrix with r rows and c columns it creates min(r-1, c) distinct principal components. Principal components are linearly uncorrelated variables created in such a way that the first principal component preserves most variance from the original dataset, with each subsequent principal component preserving less than previous one [20, 21].

#### Decision tree

Decision tree is a model, which we can imagine as a binary tree. In this tree, every node is either a decision node or the end node. Decision node contains a condition and has two child nodes. First child node represents all cases, where the condition was not met and the second child node represents all cases, where condition was met. These child nodes can be decision nodes or end nodes. End nodes do not contain any conditions and they represent the final decision made by model. Decision tree is the simplest predictive model and contains conditions exclusively. Usually, we need two sets of data to create the model, commonly called features and labels. Features represent data which are known before prediction and labels represent expected results of prediction. For example, the data representing features could be size, weight, length of nose and color and labels could symbolize if the animal described by those features is a dog or it is a cat.

#### Comparison of the Cronobacter and Citrobacter decision trees suitability for phage Dev-CS701

Table 1: Cluster number, position and predicted function of Dev-CS701 phage proteins suitable for decision trees of the genera *Cronobacter* and *Citrobacter*

| Citrobacter tree | | | Cronobacter tree | | |
|---|---|---|---|---|---|
| Position | Cluster | Present/Function | Position | Cluster | Present/Function |
| 0 | 743 | | 0 | 2261 | No |
| 1 | 170 | Hinge connector of long tail fiber distal connector | 1 | 1221 | No |
| 1 | 170 | L-shaped tail fiber protein | 2 | 247 | Yes/Clamp loader subunit |
| 2 | 918 | No | 3 | 714 | No |
| 3 | 966 | No | 4 | 426 | Yes/putative endolysin |
| 4 | 365 | No | 5 | 170 | Yes/Hinge connector of long tail fiber distal connector |
| 5 | 828 | Unknown | 5 | 170 | Yes/L-shaped tail fiber protein |
| 6 | 19 | No | 6 | 830 | Yes/Peptidoglycan binding protein |
| 7 | 186 | Portal vertex protein | 7 | 54 | Yes/MobE endonuclease |
| 8 | 23 | DNA methylase | 8 | 228 | No |
| 9 | 54 | MobE endonuclease | 9 | 640 | No |
| | | | 10 | 104 | No |
| | | | 11 | 679 | No |

### Citrobacter Decision tree

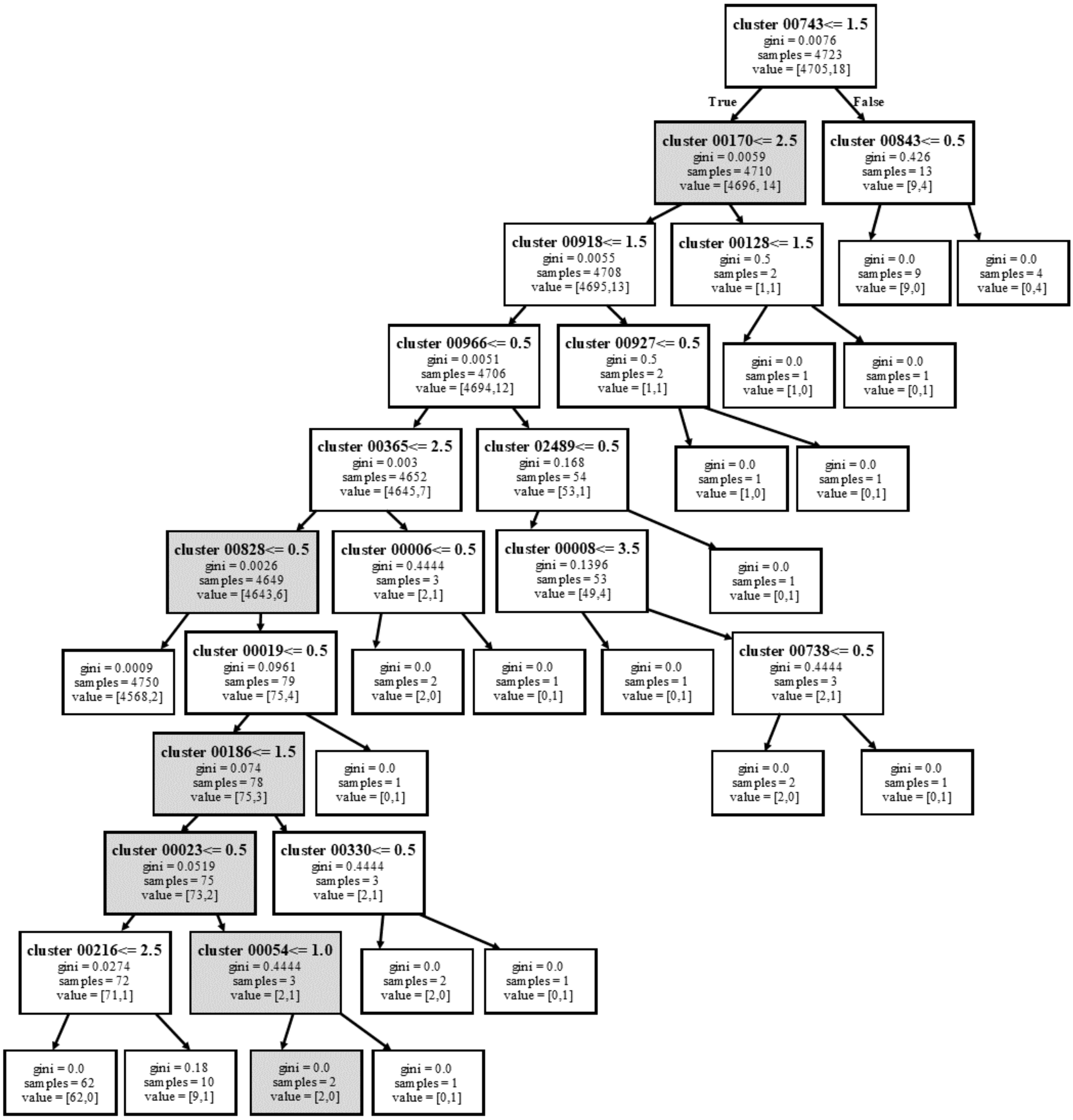


### Cronobacter decision tree

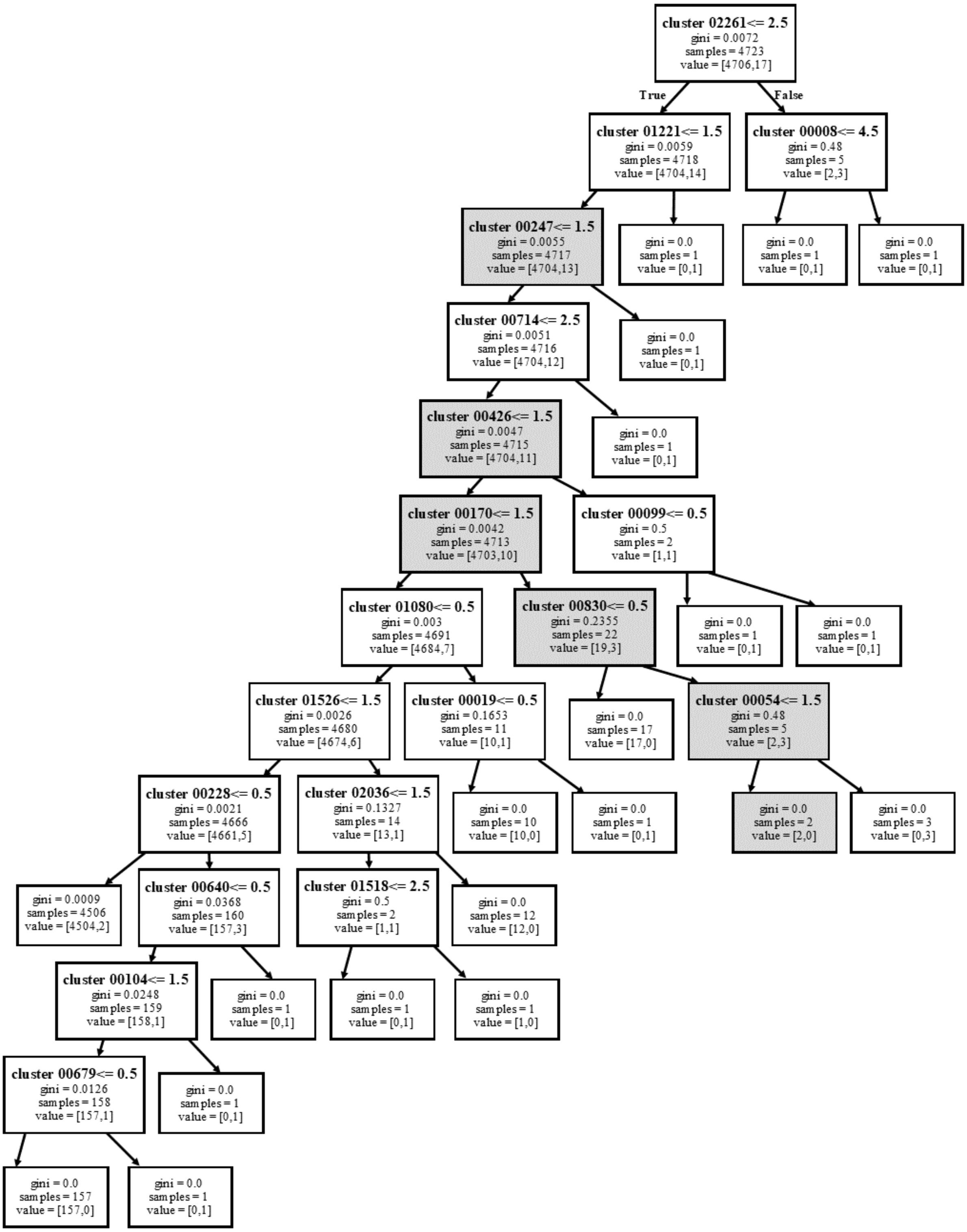


Each position in the decision tree consists of a cluster number. The cluster represents a group of proteins with a certain similarity. It further contains a condition representing the number of proteins from a given cluster that must be present in the phage being tested. The gini coefficient is a statistical measure of distribution. The coefficient ranges from 0 to 1, with 0 representing perfect equality and 1 representing perfect inequality of proteins in the cluster.

The number of samples represents how many phages meet the conditions of the previous tree levels and how this cluster divides the phages into those that meet the condition and those that do not.


1. Köster J, Rahmann S (2018) Snakemake—a scalable bioinformatics workflow engine. Bioinformatics. https://doi.org/10.1093/bioinformatics/bty350

2. Benson DA, Cavanaugh M, Clark K, et al (2018) GenBank. Nucleic Acids Res 46:D41–D47

3. Maglott D, Ostell J, Pruitt KD, Tatusova T (2011) Entrez Gene: gene-centered information at NCBI. Nucleic Acids Res 39:D52–7

4. Cock PJA, Antao T, Chang JT, et al (2009) Biopython: freely available Python tools for computational molecular biology and bioinformatics. Bioinformatics 25:1422–1423

5. Griffith M, Griffith OL (2004) RefSeq (the Reference Sequence Database). In: Dictionary of Bioinformatics and Computational Biology

14. Heringa J (2004) Needleman-Wunsch Algorithm. In: Dictionary of Bioinformatics and Computational Biology

15. Nepusz T, Sasidharan R, Paccanaro A (2010) SCPS: a fast implementation of a spectral method for detecting protein families on a genome-wide scale. BMC Bioinformatics 11:120

16. Zdobnov EM, Apweiler R (2001) InterProScan--an integration platform for the signature-recognition methods in InterPro. Bioinformatics 17:847–848

17. Hulo N (2006) The PROSITE database. Nucleic Acids Res 34:D227–D230

18. Attwood TK (2002) The PRINTS database: a resource for identification of protein families. Brief Bioinform 3:252–263

19. Corpet F (1998) The ProDom database of protein domain families. Nucleic Acids Res

26:323–326

20. Schultz J, Milpetz F, Bork P, Ponting CP (1998) SMART, a simple modular architecture research tool: Identification of signaling domains. Proceedings of the National Academy of Sciences 95:5857–5864

21. Jolliffe IT (2013) Principal Component Analysis. Springer Science & Business Media
